# Uncertainty is Maintained and Used in Working Memory

**DOI:** 10.1101/2020.10.06.328310

**Authors:** Aspen H. Yoo, Luigi Acerbi, Wei ji Ma

## Abstract

What are the contents of working memory? In both behavioral and neural computational models, a working memory representation is typically described by a single number, namely a point estimate of a stimulus. Here, we asked if people also maintain the uncertainty associated with a memory, and if people use this uncertainty in subsequent decisions. We collected data in a two-condition orientation change detection task; while both conditions measured whether people used memory uncertainty, only one required maintaining it. For each condition, we compared an optimal Bayesian observer model, in which the observer uses an accurate representation of uncertainty in their decision, to one in which the observer does not. We find that this “Use Uncertainty” model fits better for all participants in both conditions. In the first condition, this result suggests that people use uncertainty optimally in a working memory task when that uncertainty information is available at the time of decision, confirming earlier results. Critically, the results of the second condition suggest that this uncertainty information was maintained in working memory. We test model variants and find that our conclusions do not depend on our assumptions about the observer’s encoding process, inference process, or decision rule. Our results provide evidence that people have uncertainty that reflects their memory precision on an item-specific level, maintain this information over a working memory delay, and use it implicitly in a way consistent with an optimal observer. These results challenge existing computational models of working memory to update their frameworks to represent uncertainty.

## 2 Introduction

Visual working memory, the process involved in actively maintaining visual information over a short period, is essential for numerous everyday behaviors as “simple” as integrating visual information across saccades and as “complex” as reading comprehension, problem solving, and decision making (Baddeley & Hitch, 1974; Baddeley, 2003; Fukuda, Vogel, Mayr, & Awh, 2010; Conway, Kane, & Engle, 2003; Just & Carpenter, 1992). As important as it is, visual working memory is also a notoriously limited process, resulting in an imperfect and incomplete picture of the world it aims to represent.

Both behavioral (e.g., Zhang & Luck, 2008; Bays & Husain, 2008; van den Berg, Shin, Chou, George, & Ma, 2012; Fougnie, Suchow, & Alvarez, 2012) and neural (e.g., Ermentrout, 1998; Wang, 2001; Compte, 2006) models of visual working memory typically represent people’s memory as a single number, a noisy estimate of the value of the stimulus. For example, someone may remember a 34° oriented line as 37°. It is, however, important in many visual working memory decisions to represent more than just a point estimate of the remembered stimulus, but the uncertainty as well. Uncertainty is technically defined as the width of a belief distribution over a stimulus, but intuitively is a subjective measure representing how unsure an observer is about the stimulus. This is different from memory precision, which is objective and represents how precisely an observer actually remembers the stimulus. An ideal observer’s uncertainty will reflect the precision with which they remembered an item, such that they are less uncertain for more precise memories. They will use this knowledge by weighing low-uncertainty information more heavily than high-uncertainty information. In a variety of domains, this strategy would increase performance and thus should be used. For example, high uncertainty over the memory of the location of a coffee cup may result in someone looking at it before reaching for it. High uncertainty over whether a friend changed their appearance may result in someone being less likely to comment on it.

Does uncertainty get taken into account in working memory-based decisions? An intuitive first place to look is the literature on working memory confidence, since confidence can be thought of as a readout of uncertainty. Experimenters have probed memory confidence by asking people to provide a rating (Rademaker, Tredway, & Tong, 2012; Vandenbroucke et al., 2014; Samaha & Postle, 2017), choose the best remembered item (Fougnie et al., 2012; Suchow, Fougnie, & Alvarez, 2017), or make a memory-based bet (Yoo, Klyszejko, Curtis, & Ma, 2018; Honig, Ma, & Fougnie, 2020). These studies have demonstrated that people have higher working memory confidence on trials that are remembered more accurately (but see Sahar, Sidi, & Makovski, 2020; Bona, Cattaneo, Vecchi, Soto, & Silvanto, 2013; Bona & Silvanto, 2014; Vlassova, Donkin, & Pearson, 2014; Maniscalco & Lau, 2015; Adam & Vogel, 2017; Samaha, Barrett, Sheldon, LaRocque, & Postle, 2016 for conflicting results), and a computational model in which memory judgements and confidence ratings are derived from the same underlying memory precision can quantitatively account for these joint data (van den Berg, Yoo, & Ma, 2017).

All these studies ask the participant to consciously access the quality of their memory. However, in naturalistic settings, people are typically not directly interrogated about their uncertainty, but use it implicitly in order to benefit later decisions. For example, looking before reaching for one’s coffee cup or commenting on a friend’s appearance are decisions that presumably use uncertainty without conscious report. In this study, we take inspiration from perceptual decision-making studies, which have demonstrated that people implicitly incorporate uncertainty to increase behavioral performance in a variety of decision-making paradigms (e.g., van Beers, Sittig, & Gon, 1999; Ernst & Banks, 2002; Alais & Burr, 2004; Körding & Wolpert, 2004; Knill & Pouget, 2004; Ma, Navalpakkam, Beck, van den Berg, & Pouget, 2011; Jazayeri & Shadlen, 2010; Stocker & Simoncelli, 2006).

There is already some evidence that people use uncertainty implicitly in working memory-based decisions. Keshvari and colleagues had humans complete a four-item orientation change detection task (Keshvari, van den Berg, & Ma, 2012); Devkar and colleagues had humans and monkeys complete a three-item orientation change localization task (Devkar, Wright, & Ma, 2017). Stimuli in both studies were ellipses, which were independently assigned to be longer and narrower, providing “high-reliability” orientation information, or shorter and wider, providing “low-reliability” orientation information. The reliability of ellipses affected the precision with which they were encoded, and thus should have affected the memory uncertainty associated with each item. To maximize performance in both tasks, participants’ uncertainty would need to reflect this variability in item-specific precision. Both studies found that a computational model that assumes participants use item-specific uncertainty accounted better for people’s choices than alternative models.

Crucially, while these two studies provide evidence that people can implicitly use uncertainty, some experimental design choices do not allow us to conclude that people are actually maintaining uncertainty per se. First, participants in the study by Devkar and colleagues received trial-to-trial feedback on the correctness of their response. It is thus possible that participants simply learned a stimulus-response mapping (Maloney & Mamassian, 2009) rather than performing Bayesian inference or other forms of probabilistic computation (i.e., still using uncertainty in their decision; Ma, 2010). Second, precision in both studies was experimentally manipulated through ellipse reliability, which was held constant through and available after the working memory delay. Thus, participants could have used this ellipse reliability as a proxy for uncertainty (Barthelmé & Mamassian, 2010), rather than maintaining this information over the working memory delay.

Thus, the goal of this study was to investigate the conjunction of uncertainty *maintenance* and *implicit use* in a working memory task. To reach this goal, we collected data in a two-condition orientation change detection task and developed computational models to test different hypotheses about uncertainty. Intuitively, uncertainty results in a criterion shift, such that a stimulus with higher uncertainty associated with it would require a larger physical change before an observer would report that it changed. In the first condition, we established that people use uncertainty if a proxy to it is provided to them at the time of decision, replicating the results from Keshvari and others (2012). In the second condition, we asked if people still use uncertainty if this proxy is not provided at the time of decision. In other words, we asked if uncertainty is being maintained in working memory.

## 3 Experimental Methods

### 3.1 Participants

Thirteen participants (11 female; mean age *M* = 21.1 years, *SD* = 2.5) completed both conditions. All participants had normal or corrected-to-normal vision. Participants were naive to the study’s hypotheses and were paid $12/hour and a $24 completion bonus. We obtained informed, written consent from all participants. The study was in accordance with the Declaration of Helsinki and was approved by the Institutional Review Board of New York University (IRB-FY2019-2490). Seven other participants were excluded because they did not meet performance criteria (explained in the Cross-Session Procedure section).

### 3.2 Stimuli

Stimuli were four, light-grey, oriented ellipses on a medium-grey background. Each ellipse could be long or short, to provide respectively higher or lower reliability information regarding the orientation of the ellipses. All ellipses had an area of 1.19 degrees of visual angle (dva). The high-reliability ellipse had an ellipse eccentricity of 0.9, such that the major and minor axis were 1.02 and 0.37 dva, respectively. The low-reliability ellipse eccentricity was determined separately for each participant to equate performance (details in Procedure).

On every trial, a stimulus display consisted of four ellipses. The probability of each ellipse being high reliability was 0.5, independent of the reliability of the other ellipses. The location of the first ellipse was drawn from a uniform distribution between polar angles 0° and 90°. Each ellipse after that was placed such that all ellipses were 90° apart on an imaginary annulus that was 7 dva away from fixation. Afterward, the x- and y- location of the ellipses were independently jittered −0.3 to 0.3 dva. The ellipse stimuli are consistent to those in Keshavri et al.’s (2012) study. In one condition, there were additionally oriented line stimuli, which were set to have approximately the same area as the ellipses. Stimuli were displayed on a 23 inch LED monitor with a refresh rate of 60 Hz and a resolution of 1920 x 1080 pixels.

### 3.3 Procedure

#### 3.3.1 Trial Procedure

##### Ellipse condition

A trial began with a fixation cross presented for 1000 ms. Four ellipses were presented for 100 ms, followed by a 1000 ms delay, then by another four ellipses for 100 ms. On half of the trials, all ellipses in the second stimulus presentation were identical to the ellipses in the first stimulus presentation. On the other half of the trials, one ellipse changed in orientation. This change was drawn from a uniform distribution, so change of any magnitude had equal probability. Each ellipse had an equal probability of containing the change. *Change* and *no change* trials were randomly interleaved throughout the experiment. The participant indicated with a keyboard button press whether they believed there was an orientation change between the two displays. This condition is identical to the experiment done by Keshvari et al. (2012).

##### Line condition

In the Line condition, the stimuli in the second presentation were oriented lines rather than ellipses. The task was otherwise identical. An example of a trial in the Ellipse and Line conditions is illustrated in Figure 1.

**Figure 1:**
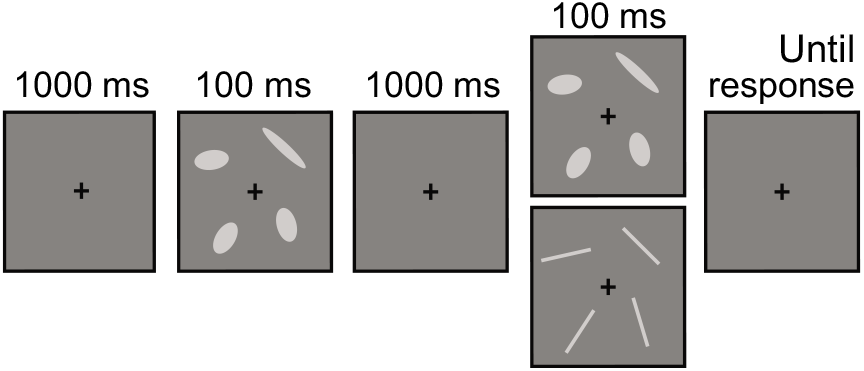
Trial sequence. Participants fixated on a cross, saw four ellipses (here showing one high-reliability ellipse and three low-reliability ellipses), maintained them over a delay, saw four stimuli again, and reported whether they believed there was an orientation change or not. In the Ellipse condition, ellipses in the second presentation were of the same reliability as in the first. In the Line condition, lines replaced ellipses in the second stimulus presentation, to avoid providing cues to the precision with which the first items were maintained.

#### 3.3.2 Cross-Session Procedure

Participants completed both conditions over six one-hour sessions. They began their first session with a Practice block, designed to ease the participants into the task. They then completed 2000 trials of each condition, preceded by a Threshold block to set the “short” ellipse reliability for each condition. Participants completed all of one condition before completing the other, and the order was counterbalanced across participants. Participants were verbally informed that each trial had a 0.5 probability of a change occurring, that a change (if present) would occur in exactly one ellipse, and the change could be “of any magnitude; big changes are as possible as small changes.” Participants were also verbally informed that some ellipses would be more elongated than others, that this may affect performance, and that half of the experiment would involve the stimuli changing from ellipses to lines. They were informed that their task did not change; the goal was always to indicate whether there was a change in orientation.

The Practice block consisted of 256 trials and was designed to ease naive participants into the speed of the task. The stimulus presentation time decreased throughout the course of the Practice block, from 333 ms to 100 ms, in 33 ms increments every 32 trials. Unlike the actual task, the ellipse eccentricities (i.e., reliabilities) of all ellipses within each trial were the same, but changed across trials. The stimuli in the second stimulus presentation corresponded to the condition that the participant completed first. For example, the stimuli in the second presentation were lines if the participant completed the Line condition first.

The Threshold block consisted of 400 trials and was used to set the ellipse eccentricity of the low-reliability ellipse in each condition. Like the Practice block, the ellipse eccentricities of all ellipses on each trial were the same, but changed on a trial-to-trial basis. The second stimulus presentation set were either ellipses or lines, corresponding to which condition the threshold was being set for. A cumulative normal psychometric function was fit to the accuracy as a function of ellipse eccentricity, and the low-reliability ellipse eccentricity was set as the value that corresponded to a predicted 65% accuracy. If the ceiling performance of the participant was estimated to be less than 75%, the Threshold block was repeated. If the psychometric function could not estimate an ellipse reliability for which performance would hit 65% after the second try, the participant was excluded from the experiment. Seven participants were excluded based on these criteria.

## 4 Experimental Results

The goal of our study was to investigate whether people maintained and used uncertainty implicitly in a working memory-based decision. To do this, we conducted a two-condition orientation change detection task. People could use memory uncertainty to maximize performance in both conditions, but only the Line condition required maintenance of that uncertainty. We conducted five repeated-measures ANOVAs to test whether condition (Ellipse, Line), the number of high-reliability ellipses displayed (*N*_high_: 0, 1, 2, 3, 4), or their interaction significantly affected the following values: proportion report “change,” false alarm rate, hit rate (for all items), hit rate (when the changed item was a low-reliability ellipse), and hit rate (when the changed item was a high-reliability ellipse). These values are visualized in Figure 2, and the statistics are reported in Table 1.

**Figure 2:**
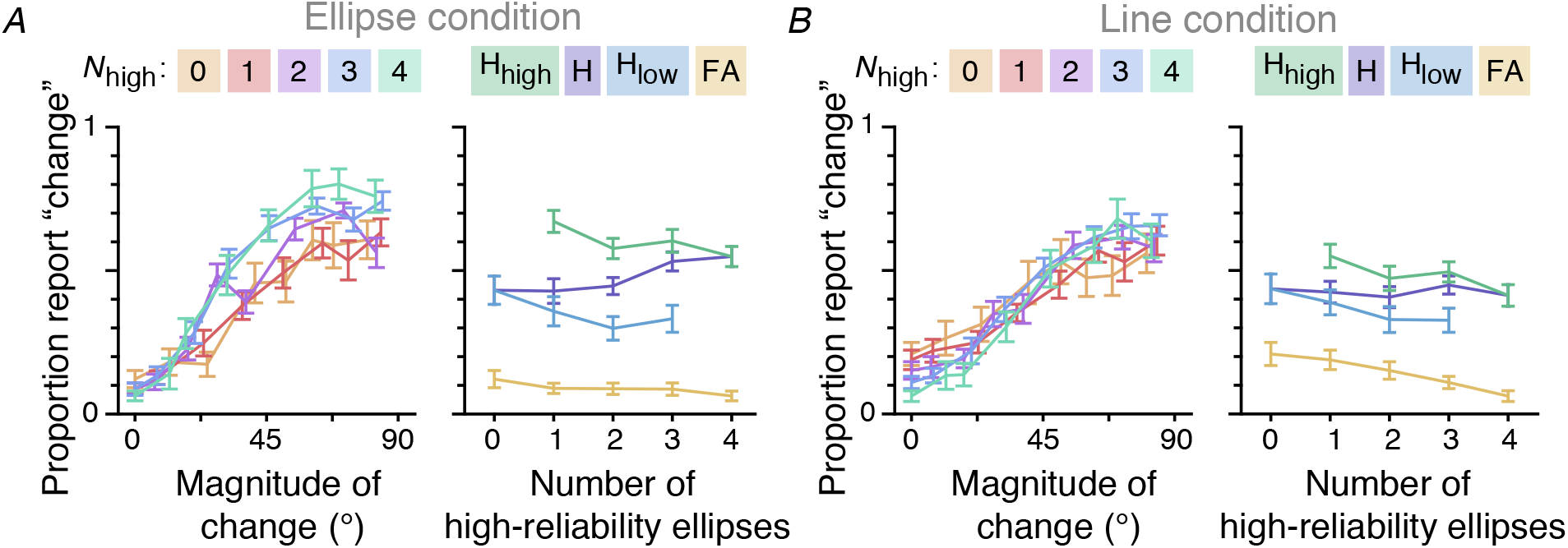
Behavioral data. Illustration of behavioral data for (*A*) Ellipse condition and (*B*) Line condition. For each condition, the left plots illustrate proportion report “change” as a function of magnitude of change. Data are binned by quantile, and different colored lines illustrate data from trials with different numbers of high-reliability ellipses presented on the first display. The right plots illustrate the proportion report “change” as a function of number of high-reliability ellipses, conditioned on whether there was no actual change (false alarm (FA): yellow), a change in a low-reliability ellipse (H_low_: blue), a change in a high-reliability ellipse (H_high_: green), or a change in any ellipse (hit (H): purple). Color legends are displayed above the plots. Note that the aggregated hits are a weighted combination of the reliability-conditioned hits. The “Z” shape formed by the hit lines are an instance of Simpson’s paradox.

**Table 1:** Results of two-way repeated-measures ANOVA. Independent variables are *N*_high_ (0,1,2,3,4) and condition (Ellipse, Line), and dependent variables are displayed as the first column. Statistics of significant effects are bolded. For all ANOVAs, we report the Greenhouse-Geisser corrected results and ε (sphericity correction) when appropriate.

| Dependent variable | Factor | Statistics | $p$ | $\epsilon$ | $\eta^2$ |
| --- | --- | --- | --- | --- | --- |
| Proportion report “change” | $N_{\text{high}}$ | $F(1.38, 16.50) = 3.37$ | 0.07 | 0.34 | 0.03 |
| | Condition | $F(1, 12) = 1.33$ | 0.27 | – | 0.01 |
| | $N_{\text{high}}$ x condition | **$F(2.12, 25.38) = 6.32$** | **0.005** | **0.52** | **0.04** |
| False alarm rate | $N_{\text{high}}$ | **$F(1.93, 23.17) = 18.21$** | **$2.07 \times 10^{-5}$** | **0.48** | **0.14** |
|  | Condition | **$F(1, 12) = 6.50$** | **0.03** | – | **0.08** |
| | $N_{\text{high}}$ x condition | **$F(1.95, 23.36) = 4.94$** | **0.02** | **0.49** | **0.05** |
| Hit rate (all) | $N_{\text{high}}$ | **$F(1.36, 16.30) = 5.29$** | **0.03** | **0.34** | **0.04** |
| | Condition | $F(1, 12) = 2.47$ | 0.14 | – | 0.03 |
| | $N_{\text{high}}$ x condition | **$F(2.04, 24.48) = 5.33$** | **0.01** | **0.51** | **0.03** |
| Hit rate (low-reliability) | $N_{\text{high}}$ | **$F(1.76, 21.07) = 23.26$** | **$8.43 \times 10^{-6}$** | **0.59** | **0.08** |
| | Condition | $F(1, 12) = 0.29$ | 0.60 | – | 0.005 |
| | $N_{\text{high}}$ x condition | $F(2.01, 24.15) = 0.37$ | 0.69 | 0.67 | 0.002 |
| Hit rate (high-reliability) | $N_{\text{high}}$ | **$F(1.98, 23.79) = 35.44$** | **$7.72 \times 10^{-8}$** | **0.66** | **0.13** |
|  | Condition | **$F(1, 12) = 14.66$** | **0.002** | – | **0.17** |
| | $N_{\text{high}}$ x condition | $F(2.15, 25.80) = 0.75$ | 0.49 | 0.72 | 0.003 |

There was a statistically significant interaction between *N*_high_ and condition on proportion report “change.” In only the Ellipse condition, the proportion report “change” was modulated by the number of high-reliability ellipses (left plot of Fig. 2 A, B). There were significantly more false alarms in the Line condition (*M* = 0.14, *SEM* = 0.03) than in the Ellipse condition (*M* = 0.09, *SEM* = 0.02; yellow lines in right plots of Fig. 2 A, B). Perhaps people confused changes in stimuli as changes in orientation.

Both reliability-conditioned hit rates (blue and green lines in right plots of Fig. 2 A, B) as well as false alarm rates decreased with increasing *N*_high_. Additionally, participants had significantly lower high-reliability hits in the Line condition and the Ellipse condition. These results could be potentially explained by participants using uncertainty information. As the total number of high-reliability ellipses, *N*_high_, increases, the number of high-reliability ellipses that do not change also increases. If people weigh high-reliability information more heavily than low-reliability information, then as the amount of high-reliability “no change” information increases, the proportion that the participants respond “change” should decrease. This would result in a decrease in reliability-conditioned hit rates and false alarm rates with increasing *N*_high_.

There is an interesting reverse in the qualitative trend when looking at all hit rates across all trials: hit rate increases as a function of *N*_high_. This Simpson’s paradox is a result of weighted averaging and the performance difference between the reliability-conditioned hit rates. As the number of high-reliability ellipses in a display increases, so does the probability of a change occurring in a high-reliability ellipse. Thus, the total hit rates for higher *N*_high_s contain more high-reliability hits than low-reliability hits, driving this value upward. Similarly, the trials to compute hit rates for lower *N*_high_s predominantly contain changes in low-reliability ellipses, thus driving the average downward. There was also a significant effect of condition; hit rates were higher in the Ellipse condition.

These statistics show that differences between factors and conditions exist, but are dissatisfying because they do not offer explanations of what these differences mean. In fact, the intuitions presented in this section are largely driven by knowledge about how noise affects decisions, knowledge acquired from computational models like signal detection theory (e.g., Green & Swets, 1966) and Bayesian decision theory. Thus, in this paper we directly test our intuitions about the underlying working memory processes through computational modeling. Computational modeling allows us to make explicit assumptions and precise quantitative predictions, which provide committal, falsifiable explanations of the processes involved.

## 5 Modeling Methods

To test whether people are maintaining and using uncertainty when making their change detection decision, we use Bayesian observer models (Ma, 2019). Bayesian models provide a normative, flexible, and interpretable framework to study the working memory process. These models are particularly useful in cases where the observer is trying to make a decision without full knowledge of task-relevant information. In working memory, people do not have full knowledge because information is not remembered perfectly. While Bayesian decision theory describes how an observer should behave in order to maximize performance, different components of the model can be easily substituted with incorrect beliefs or suboptimal use of information, and thus provides a good template for building models with “imperfectly optimal observers” (Maloney & Zhang, 2010) or “model mismatch” (Orhan & Jacobs, 2014; Beck, Ma, Pitkow, Latham, & Pouget, 2012; Acerbi, Vijayakumar, & Wolpert, 2014).

We model the observer’s decision process as consisting of an encoding stage and a decision stage. The encoding stage describes the task statistics and our assumptions about how memories are generated. In the decision stage, the observer calculates a decision variable based on their belief of the encoding stage and decides whether to report “change” or “no change” based on some decision rule. We compared two models: one in which uncertainty is maintained and used and another that is not, named the “Use Uncertainty” and the “Ignore Uncertainty” model, respectively. This section describes how these models were defined, fit, and compared.

### 5.1 Encoding Stage

In this section, we define the statistical structure of the experiment and define our assumptions about how memories are generated in an observer. On every trial, there is a 0.5 probability of there being a change, *p*(*C* = 1) = 0.5, where *C* takes values 0 (no change) and 1 (change). On change trials, exactly one item changes in its orientation, and each item is equally probable to be changed. The orientation change, Δ, is drawn from a uniform distribution, 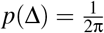. (For mathematical convenience, and without loss of generality, we doubled the actual orientation of stimuli in all model specifications such that the values span 0 to 2π rather than 0 to π. We do not double these values when illustrating model fits.)

We denote the vector of all orientations of the items presented on the first display by **ξ**, in which each element is an independent draw from a uniform distribution over orientation space. The vector of orientations at the second display, **ϕ**, was identical to **ξ** in no change trials. In change trials, the *i*th element of **ϕ**, the location of change, was equivalent to ξ*_i_* + Δ.

We model the memory process for each item of each display according to the Variable Precision model (van den Berg et al., 2012; Fougnie et al., 2012), by which memories are described as a continuous resource that randomly fluctuates across items and trials. The noisy measurements of each item on each display, ***x*** = (*x*_1_,…,*x_N_*) and ***y*** = (*y*_1_,…,*y_N_*), are conditionally independent and drawn from a Von Mises distribution centered on the actual orientation presentation,

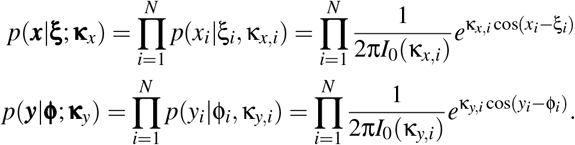

The κs are the concentration parameter of the Von Mises distribution, and are related to the precision with which each item is remembered; a higher κ corresponds to higher precision. The subscript of each κ indicates which item it refers to (e.g., κ_*x,i*_ is concentration parameter for *x_i_*, the *i*^th^ item the first stimulus presentation). We assume that memory precision varies across items, above and beyond the precision differences due to stimulus reliability. In other words, κ_*x,i*_ and κ*_y,i_* are themselves random variables, rather than single values. Rather than sampling κ itself, we sample the Fisher information of the Von Mises distribution, *J*, from a gamma distribution:

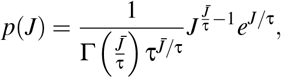

where τ is the scale parameter of the gamma distribution and 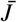 is the mean precision. The relationship between *J* and κ is the following:

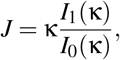

where *I*_0_ is a modified Bessel function of the first kind of order 0 and *I*_1_ is a modified Bessel function of the first kind of order 1 (van den Berg et al., 2012; Keshvari et al., 2012). We allow the mean precision to differ across stimulus shape; the precisions of memories corresponding to low-reliability ellipses are drawn from a gamma distribution with mean 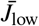 and high-reliability ellipses with mean 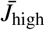. Parameter τ is shared across both distributions. Because items in the first display were presented earlier, there are certainly differences in the precision with which items in the first and second display are maintained, independent of ellipse reliability. However, the amount that the first and second displays contribute to the overall measured change are extremely hard to tease apart in the model. Thus, we use one parameter per reliability and recognize that this estimate will be some average of the precisions of the first and second display.

When modeling the Line condition, we have an additional parameter, 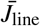, which corresponds to the mean precision with which each line on the second display is remembered by the observer. To limit model complexity, the gamma function from which each line’s precision is drawn shares the same scale parameter τ as the distributions from which the ellipse precisions are drawn.

### 5.2 Decoding Stage

#### 5.2.1 Decision Variable

The essence of Bayesian inference is that an observer can compute a posterior over task-relevant latent variables, and should if they want to maximize performance. In this case, the observer should calculate the probability of the state of the world (i.e., change or no change) given their observations, *p*(*C*|***x***, ***y***), which they can compute using Bayes rule. With a scenario in which there are only two states of the world, it is convenient to combine these into a ratio. Thus, we assume the observer calculates, for each item, the ratio of the likelihood of there being change and the likelihood of there being no change:

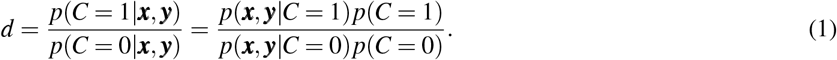

Details of the derivation can be found in Appendix 8.1, but this simplifies to the following expression:

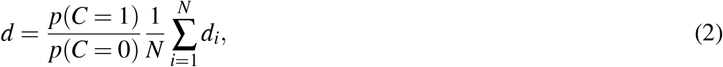

where

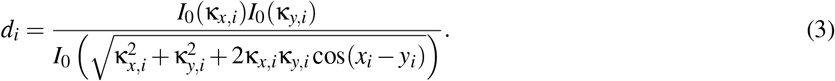

*I*_0_ is a modified Bessel function of the first kind of order 0, and the κs are the concentration parameters of the noise distributions for the item indicated in the subscript. Intuitively, *d_i_* provides a measure of the evidence of change for the *i*^th^ item. It increases with the measured amount of change, *x_i_*, – *y_i_*, weighted by a function of the precisions with which *x_i_* and *y_i_* are remembered. The *d_i_*s are averaged in the decision variable *d*, providing the optimal measure of evidence of change of the entire display.

This is the step in which the use of uncertainty comes in. Observers who correctly maintain and use uncertainty (i.e., observers who act in accordance with the optimal, “Use Uncertainty” model) compute *d_i_* exactly as described. However, observers acting in accordance with the “Ignore Uncertainty” model do not know or do not consider that the precision of their memories for all items in both displays varies. Computing the decision rule for the Ignore Uncertainty observer is the same as replacing all κs in Eq. 3 with a constant, resulting in the following local decision variable:

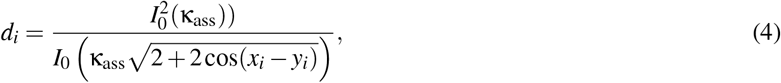

where κ_ass_ is the assumed precision for all items on all displays. The decision variable thus becomes just a function of cos(*x_i_* – *y_i_*), because the remainder of the expression is constant. The Ignore Uncertainty observer thus ignores any factor that could have affected their memory precision. We recognize this is a strong assumption, and we weaken it in the subsequent Model Variants section.

#### 5.2.2 Decision Rule

The observer maps this decision variable onto a response by reporting “change” whenever the probability of there being a change is greater than 0.5 (Figure 3). An optimal observer would thus report “change” when the ratio of the likelihood of there being a change and the likelihood of there being no change (Eq. 2) is greater than 1. For convenience, we use the logarithm of the likelihood ratio; the optimal observer would thus report “change” if this value is greater than 0. However, we allow the observer to have some response bias (e.g., due to unequal priors, rewards, or motor costs), and thus implement the following decision rule:

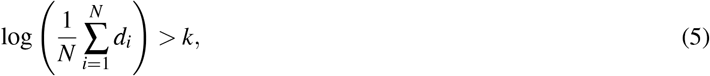

where *k* is a free parameter. For both models, we implemented global decision noise by adding zero-mean Gaussian noise with standard deviation σ_d_ to the log of decision variable *d* (Keshvari et al., 2012; Acerbi et al., 2014; Mueller & Weidemann, 2008). Additionally, participants randomly guess with probability λ, due to factors such as lapses in attention.

**Figure 3:**
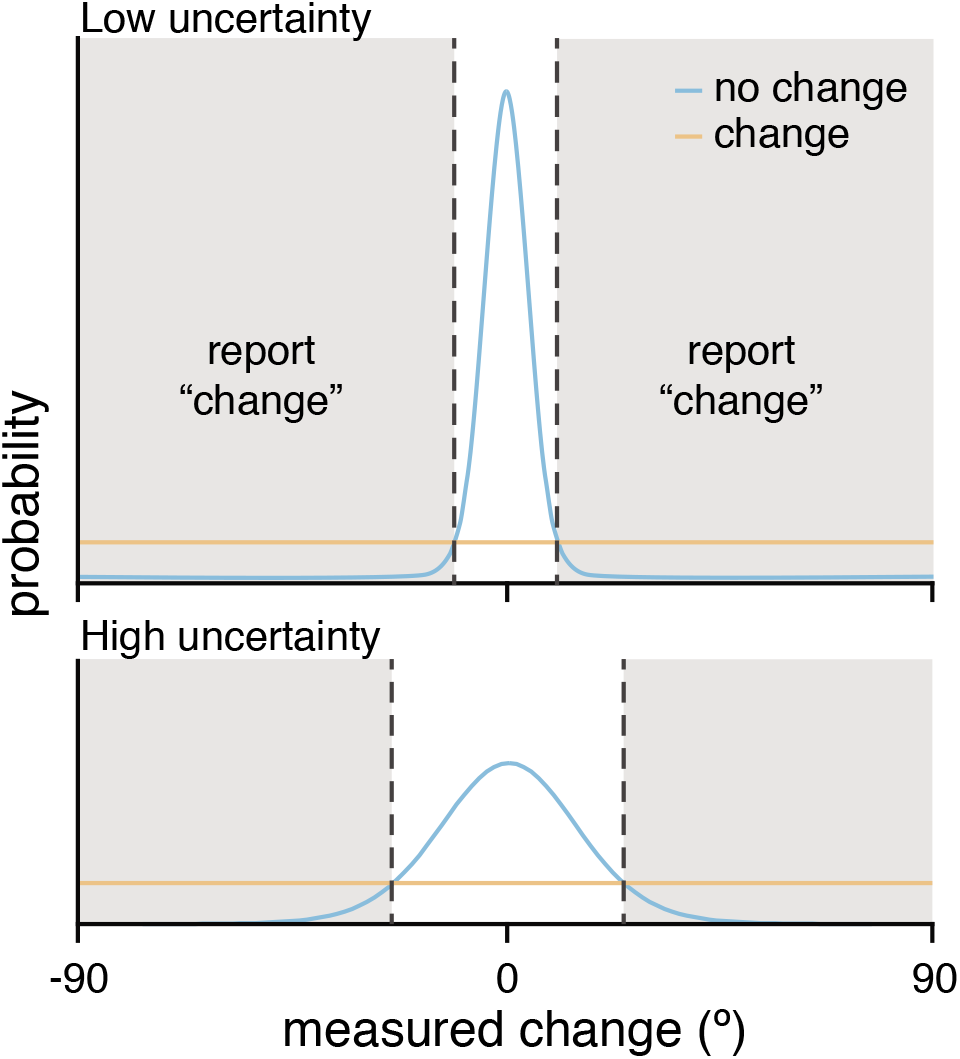
Model didactics. This didactic illustrates a simplified one-item version of this task. The probability of the measured change for an item given that the item did (orange) or did not (blue) change orientation, as estimated by the optimal observer. Uncertainty modulates the width of the no change distribution, such that higher uncertainty makes the no change distribution wider (bottom). The optimal observer (with *k* = 0) places their decision boundaries at the intersection of the change and no change distributions (vertical dashed lines), reporting “change” whenever that state of the world is more probable (shaded region) and “no change” otherwise.

### 5.3 Parameter Estimation and Model Comparison

#### 5.3.1 Parameters

Both models in both conditions have parameters 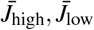, τ, *k*, λ, and σ_d_. Parameters 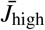 and 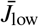 correspond to the mean precision of the high- and low-reliability ellipses, respectively. Precision is also affected by the scale parameter of the gamma distribution from which item-wise precision is drawn, τ; this value is shared across the two ellipse types and the line when applicable. Parameter *k* is the observer’s response bias; λ is the probability on each trial that the observer lapses and responds randomly; σ_d_ is the standard deviation of the Gaussian from which decision noise is simulated.

When fitting data from the Line condition, there is an additional parameter 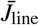, corresponding to the mean precision with which the line stimulus is represented. The Ignore Uncertainty model has one additional parameter: *J*_ass_, the assumed precision of all stimuli in both displays.

#### 5.3.2 Parameter Estimation

The likelihood of the parameter combination **θ** for a given participant and model is the probability of the data given the parameter combination. We used the log likelihood, which we denote LL:

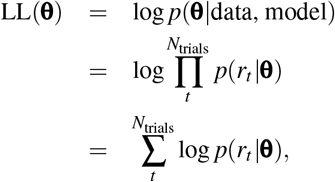

where *r_t_* is the participant’s response on the *t*^th^ trial. For each participant, we used maximum-likelihood estimation to find which parameter combination best describes participant’s data. Computing the LL analytically is intractable, so we used Inverse Binomial Sampling (IBS; van Opheusden, Acerbi, & Ma, 2020), a method which efficiently computes an unbiased estimate of the LL. This calculation is stochastic, so we additionally used an optimization algorithm that can account for stochasticity and expensive LL evaluations (BADS; Acerbi & Ma, 2017). BADS explicitly incorporates uncertainty in the estimated LL and converges in fewer function evaluations than other stochastic optimization methods (e.g., CMA-ES, genetic algorithms), making it an ideal optimization method when likelihood calculations are computationally expensive and stochastic. We used 20 different starting positions, using Latin hypercube sampling, to reduce the probability of finding a local minimum. We took the parameter combination corresponding to the minimum negative log-likelihood of our runs as the ML parameter estimate. The estimated LL at the candidate optimum was reevaluated using 1000 repetitions in IBS, in order to reduce the standard deviation of estimation noise to less than 1. We denote the maximum log-likelihood by LL*.

#### 5.3.3 Model Comparison

We compared models using corrected Akaike Information Criterion (AICc; Hurvich & Tsai, 1987) and the Bayesian Information Criterion (BIC; Schwarz, 1978). BIC penalizes for number of model parameters *N*_pars_ harsher than AICc does.

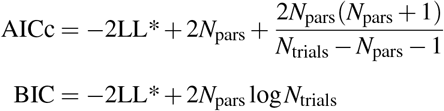

### 5.4 Modeling Results

We compared the fits of the Use Uncertainty and Ignore Uncertainty models to each of the conditions separately. The Use Uncertainty model provides a good qualitative fit to the data in both conditions (top row of Figure 4A), while the Ignore Uncertainty model is unable to capture the data (bottom row of Figure 4A). To compare models quantitatively, we used summed ΔAICc and ΔBIC, that is the difference of summed AICc (resp. BIC) across participants between the Ignore Uncertainty and Use Uncertainty models (positive values mean that Use Uncertainty fits better). Summing model comparison metrics across participants implicitly assumes that all participants are fit by the same model. For both conditions and model comparison metrics, participants were better fit by the Use Uncertainty model than the Ignore Uncertainty model (summed [95% bootstrapped confidence interval (CI)] ΔAICc across subjects – Ellipse: 3091 [2015, 4321], Line: 2764 [1468, 4400]. ΔBIC – Ellipse: 3263 [2155, 4450], Line: 2935 [1640, 4433]). Note that, while reporting the summed ΔAICc and ΔBIC, for visualization we plot the individual differences and plot the 95% CIs of the median ΔAICc (Figure 4B). Parameter estimates for the Use Uncertainty model in the Ellipse and Line condition can be found in the Appendix (Tables 4 and 5, respectively).

**Figure 4:**
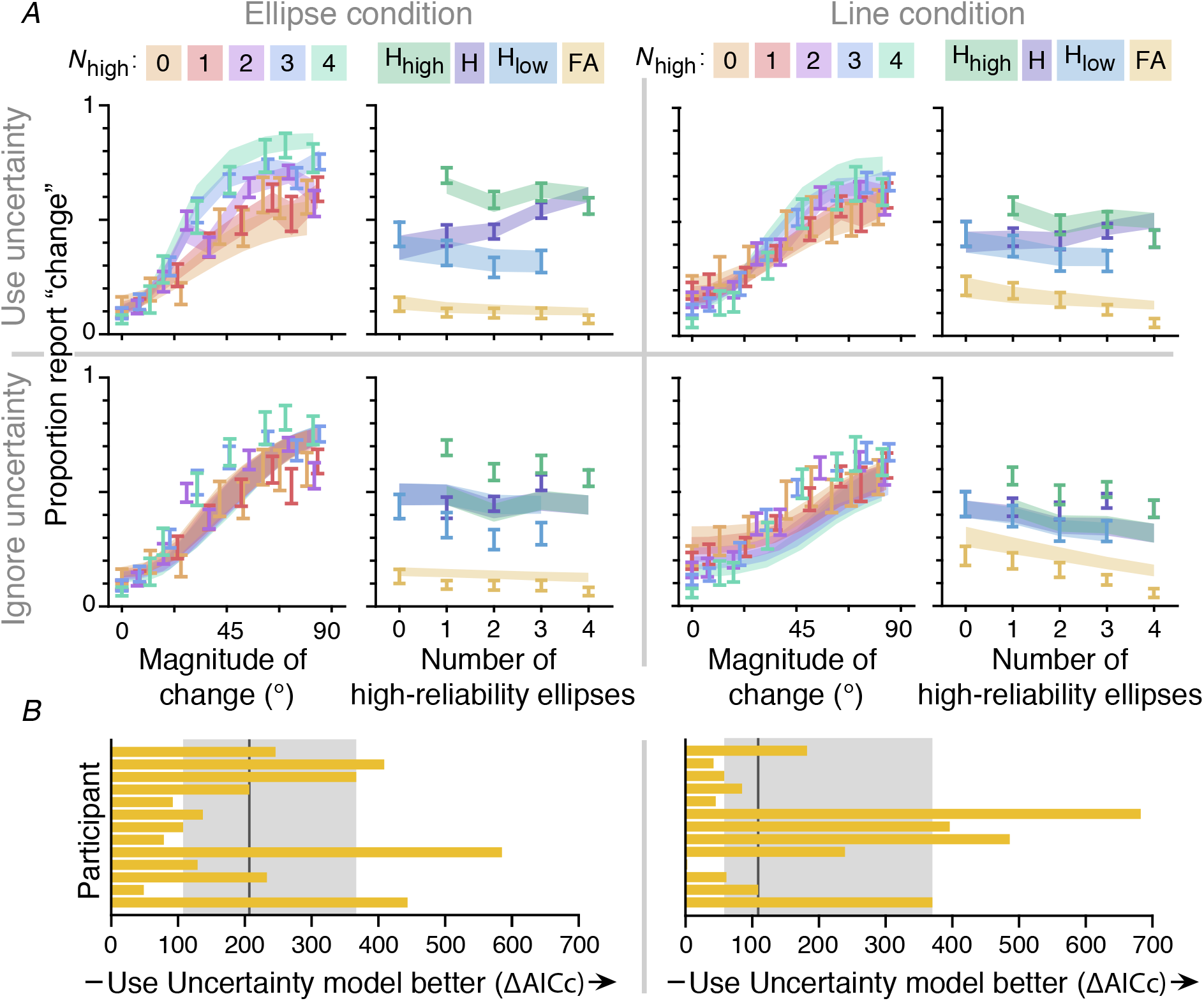
Model fits. *A*. *M*±SEM data (error bars) and model fits (fills) for the Use (top) and Ignore (bottom) Uncertainty models and the Ellipse (left) and Line (right) conditions. For each model and condition, the left graph illustrates the proportion report “change” as a function of amount of change. Data and models are binned by quantiles, and color indicates the number of high-reliability ellipses in the first display. The right graph illustrates the proportion hits for high-reliability items (green), hits for low-reliability items (blue), total hits (purple), and false alarms (yellow) as a function of number of high-reliability items. *B*. Model comparison for the Ellipse (left) and Line (right) conditions. Each bar indicates the individual-subject ΔAICc between the Use and Ignore Uncertainty models, where a positive value indicates that the Use Uncertainty model is favored. The vertical grey line indicates the median across participants, and the shaded region illustrates the 95% bootstrapped confidence interval of the median. Only ΔAICcs are illustrated here; ΔBICs gave similar results.

## 6 Model Variants

While the Use Uncertainty model provides a good fit to the data, the two models we have considered thus far contain strong assumptions that uncertainty is either perfectly used or entirely ignored. In this section, we modify the assumptions by factorially comparing different formulations of the encoding, inference, and decision stage of the model (van den Berg, Awh, & Ma, 2014; Acerbi, Wolpert, & Vijayakumar, 2012; Keshvari et al., 2012). A factorial model comparison is an effective way of testing which of the assumptions we made were critical for accounting for human behavior, and thus which are reasonable to make conclusions about. In this section, we demonstrate that our general conclusions about the use of uncertainty do not depend on the specific assumptions we made when defining our model. We only discuss the results of the Line condition here, since it is the only condition that investigates the maintenance of uncertainty in working memory. However, we did the same analysis to the Ellipse condition data and found consistent results (Appendix 8.3).

### 6.1 Encoding

In both the Use and Ignore Uncertainty models, we assumed that observers’ encoding noise followed that of a Variable Precision model (van den Berg et al., 2012; Fougnie et al., 2012). Here, we also consider that observers’ memory precision varies only based on stimulus type, and does not fluctuate on an item-to-item basis. With this “Fixed Precision” assumption of encoding noise, the κ for each item is determined only by its stimulus type; high-reliability ellipses would be encoded with parameter κ_high_, low-reliability ellipses with κ_low_, and lines with κ_line_.

### 6.2 Inference

Observers calculate the decision variable according to some inference process, which we allow to be independent of the true generative process. The potential model mismatch (Orhan & Jacobs, 2014; Beck et al., 2012; Acerbi et al., 2014) between the true and believed generative process could be due to a result of wrong beliefs about the generative process or computation limitations that prevent accurate representation of the generative model. We consider that observers may use partial knowledge of uncertainty, rather than fully Using or Ignoring uncertainty.

We consider that the observer may have one of four inference models, listed below in decreasing order of how many factors the observer takes into account in their uncertainty:

1. Variable precision (V): the observer believes that mean memory precision varies with the exact stimulus shape (low-reliability ellipse, high-reliability ellipse, line) and that there is additional noise for each item at each presentation. This inference model is optimal when the true generative process is Variable precision.
2. Fixed precision (F): the observer believes that memory precision varies with the exact stimulus shape (low-reliability ellipse, high-reliability ellipse, line), but does not consider that there is additional noise for each item at each presentation. This inference model is suboptimal when the true generative process is Variable precision, but optimal when the true generative process is Fixed precision.
3. Limited (L): the observer believes that memory precision varies across shapes (ellipse vs. line). This observer does not consider differences in precision between high- and low-reliability ellipses or additional noise for each item at each presentation. This observer is suboptimal.
4. Same precision (S): the observer believes that memory precision is the same throughout the condition, and does not vary with stimulus shape or anything else. This is the “Ignore Uncertainty” observer and is suboptimal.

Note that the Variable and Same precision inference schemes here are identical to that of Keshvari and others’ (2012), and the Fixed precision here is equivalent to their “Equal” precision inference scheme.

### 6.3 Decision Rule

The Use and Ignore Uncertainty models use the optimal decision rule (Eq. 5). Note that participants may have incorrect assumptions about the noise in their memory, but still be acting in accordance with Bayesian decision theory (i.e., still using the correct decision rule), resulting in “imperfectly optimal observers” (Maloney & Zhang, 2010). Alternatively, participants could be calculating the optimal decision variable, but be using a suboptimal decision rule. Here, we consider observers who use the max rule, reporting “change” whenever the maximum evidence of change is greater than some criterion, *k*,

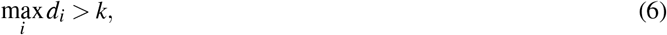

rather than averaging *d_i_*s. These observers are not Bayes-optimal, but are still using probabilistic computation (i.e., still using their uncertainty) in the calculation of *d_i_*. In fact, in many cases these decision rules do not result in substantially different behavior (Ma, Shen, Dziugaite, & van den Berg, 2015). For example, if all *d_i_*s are similar, then a max and an average will result in similar values. If the maximum *d_i_* is substantially larger than the others, both decision rules can result in similar behavior by adjusting *k*.

### 6.4 Parameters

There are two possible encoding schemes ((V)ariable, (F)ixed), four possible inference schemes ((V)ariable, (F)ixed, (L)imited, (S)ame), and two possible decision rules ((O)ptimal, (M)ax). Factorially combining each of these characteristics would yield 16 different models. We choose not to consider the models in which the generative model is “F” but the observer assumes “V” under the assumption that people tend not to assume the (perceptual) world is more complicated than it actually is; thus, we test a total of 14 models. We denote each model by the letters corresponding to their encoding scheme, inference scheme, and decision rule (e.g., VVO is the model with Variable precision encoding, an observer assumes Variable precision, and an Optimal decision rule). The VVO model is the Use Uncertainty model; the VSO model is the Ignore Uncertainty model.

#### Encoding parameters

Like before, observers with Variable precision encoding have parameters 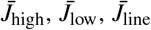, and τ. Observers with Fixed precision encoding have parameters *J*_high_, *J*_low_, and *J*_line_.

#### Inference parameters

For the observer who correctly infers their encoding process (i.e., VVO, VVM, FFO, or FFM), there are no additional parameters. If the observer has Variable precision encoding but does not take into account individual-item variations (i.e., VFO or VFM), then the assumed precision is 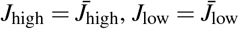, and 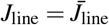 for high-reliability ellipses, low-reliability ellipses, and lines, respectively. Limited inference observers (i.e., VLO, VLM, FLO, FLM) have two additional parameters: *J*_ass, e_ and *J*_ass, 1_, corresponding to the assumed precision of the ellipses and lines, respectively. Same inference observers, who do not take any memory variations into account (i.e., VSO, VSM, FSO, FSM), have one additional parameter *J*_ass_, corresponding to the assumed precision of all items.

#### Decision parameters

Observers using both the optimal or max decision rule have parameter *k*, corresponding to the decision criterion. If any item has a decision variable greater than *k*, then they will report “change.”

Each model and their corresponding parameters is listed in Table 2. Note that the Same inference observer who uses the max rule (i.e., VSM, FSM) has one less parameter than their Optimal decision rule counterpart (i.e., VSO, FSO) because making a decision depends only on the item with the largest measured change.

**Table 2:** Model parameters. Model parameters for Line condition. Parameters **not used** for fitting the Ellipse condition are displayed in parentheses. The top colored cell corresponds to parameters of the Use Uncertainty (VVO) model. The bottom colored cell corresponds to the parameters of the Ignore Uncertainty (VSO) model.

| Encoding | Inference | Decision Rule |  |
| --- | --- | --- | --- |
|  |  | (O)ptimal | (M)ax |
| (V)variable | (V)variable | $\bar{J}_{\text{high}}, \bar{J}_{\text{low}}, \tau, k, \lambda, \sigma_d(\bar{J}_{\text{line}})$ | $\bar{J}_{\text{high}}, \bar{J}_{\text{low}}, \tau, k, \lambda, \sigma_d(\bar{J}_{\text{line}})$ |
| | (F)ixed | $\bar{J}_{\text{high}}, \bar{J}_{\text{low}}, \tau, k, \lambda, \sigma_d(\bar{J}_{\text{line}})$ | $\bar{J}_{\text{high}}, \bar{J}_{\text{low}}, \tau, k, \lambda, \sigma_d(\bar{J}_{\text{line}})$ |
| | (L)imited | $\bar{J}_{\text{high}}, \bar{J}_{\text{low}}, \bar{J}_{\text{ass}, e} \tau, k, \lambda, \sigma_d(\bar{J}_{\text{line}}, \bar{J}_{\text{ass}, l})$ | $\bar{J}_{\text{high}}, \bar{J}_{\text{low}}, \bar{J}_{\text{ass}, e} \tau, k, \lambda, \sigma_d(\bar{J}_{\text{line}}, \bar{J}_{\text{ass}, l})$ |
| | (S)ame | $\bar{J}_{\text{high}}, \bar{J}_{\text{low}}, \bar{J}_{\text{ass}}, \tau, k, \lambda, \sigma_d(\bar{J}_{\text{line}})$ | $\bar{J}_{\text{high}}, \bar{J}_{\text{low}}, \tau, k, \lambda, \sigma_d(\bar{J}_{\text{line}})$ |
| (F)ixed | (F)ixed | $J_{\text{high}}, J_{\text{low}}, k, \lambda, \sigma_d(J_{\text{line}})$ | $J_{\text{high}}, J_{\text{low}}, k, \lambda, \sigma_d(J_{\text{line}})$ |
| | (L)imited | $J_{\text{high}}, J_{\text{low}}, J_{\text{ass}, e}, k, \lambda, \sigma_d(J_{\text{line}}, J_{\text{ass}, l})$ | $J_{\text{high}}, J_{\text{low}}, J_{\text{ass}, e}, k, \lambda, \sigma_d(J_{\text{line}}, J_{\text{ass}, l})$ |
| | (S)ame | $J_{\text{high}}, J_{\text{low}}, J_{\text{ass}}, k, \lambda, \sigma_d(J_{\text{line}})$ | $J_{\text{high}}, J_{\text{low}}, k, \lambda, \sigma_d(J_{\text{line}})$ |

### 6.5 Model Comparison Results

#### 6.5.1 Comparison of individual models

As previously described, we estimated parameters for each participant and compared models using AICc and BIC. In this section, we only discuss the results of the Line condition using summed AICc and BIC differences between the VVO (Use Uncertainty) and other models. We only discuss the data from the Line condition because it is the only condition that allows us to interrogate whether people are *maintaining* the uncertainty that they use in the change detection decision. For completeness, we report the results of the Ellipse condition in the Appendix 8.3.

When using AICc, the VVO model seems to be able to capture the human data the best, indicated by a positive summed ΔAICc and 95% bootstrapped CIs compared to all alternative models. When using ΔBIC, the VVO model still fits best, but the 95% CIs are not above 0 for the FFO model, indicating that VVO does not fit the data significantly better than the FFO model. Qualitatively, both VVO and FFO models fit the data well (Figure 5). Note that, while reporting the summed ΔAICc and ΔBIC, we plot the individual differences and plot the 95% bootstrapped CIs of the median ΔAICc.

**Figure 5:**
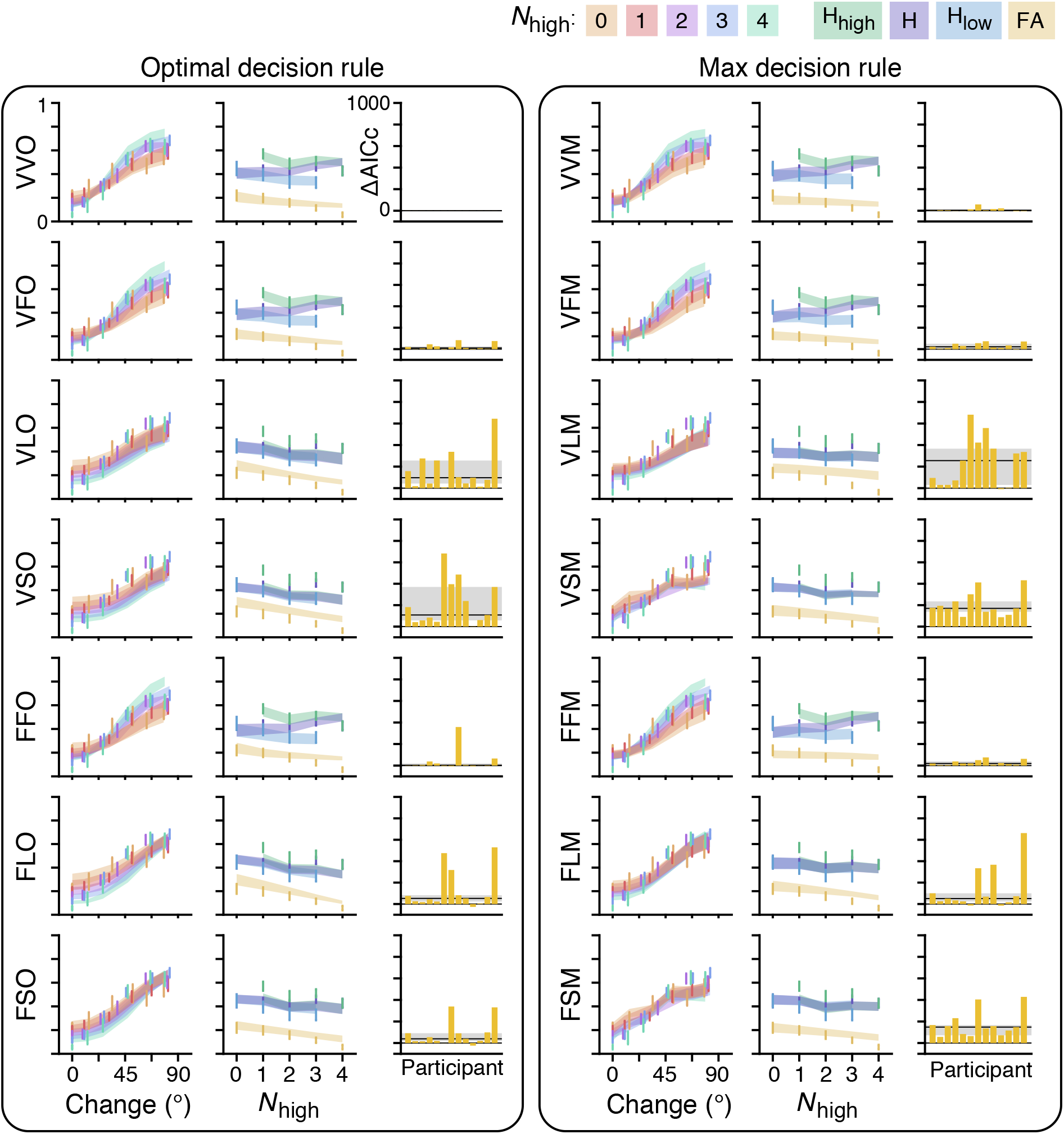
Factorial model comparison. Model predictions and performance of all possible combinations of different encoding, inference, and decision rules. *M* ± *SEM* data (error bars) and model fits (fills) for all models, organized into two columns by decision rule. For each model (each row within each column), the left graph illustrates the proportion report “change” as a function of amount of change. Color indicates the number of high-reliability ellipses (legend at the top of the figure). The middle graph illustrates the proportion hits for high-reliability items (green), hits for low-reliability items (blue), hits averaged across the display (purple), and false alarms (yellow) as a function of number of high-reliability items (legend at the top right of the figure). The right graph illustrates the individual-participant ΔAICc, where positive numbers indicate the VVO model is a better fit to the data. The grey horizontal line and shaded region illustrates median and the 95% bootstrapped confidence interval of the median across participants.

In the Appendix, we additionally report the results of group Bayesian Model Selection (BMS) for both conditions (Appendix 8.4). While summing the ΔAICc and ΔBIC implicitly assumes that participants are all fit by the same model, group BMS allows for participant heterogeneity and directly infers the distribution of participants across models. Using this alternative model comparison metric does not really change our results; VVO and FFO fit the participants’ data substantially better than other models, but their performance against one another depend on the model comparison metric.

**Table 3:** Summed ΔAICc and ΔBIC: Line condition. The sum and 95% bootstrapped confidence interval of the AICc and BIC differences between the optimal VVO (Use Uncertainty) model and others. A positive value indicates that the VVO model provides a better fit to the data. The cells corresponding to the Use (VVO) and Ignore (VSO) Uncertainty models are colored in blue.

| Encoding | Inference | Decision Rule |  |  |  |
| --- | --- | --- | --- | --- | --- |
|  |  | (O)ptimal |  | (M)ax |  |
| | | $\Delta\text{AICc}$ | $\Delta\text{BIC}$ | $\Delta\text{AICc}$ | $\Delta\text{BIC}$ |
| (V)variable | (V)variable | 0 [0, 0] | 0 [0, 0] | 119 [19, 247] | 119 [26, 247] |
|  | (F)ixed | 295 [119, 502] | 295 [114, 477] | 381 [201, 569] | 381 [200, 578] |
|  | (L)imited | 2069 [957, 3433] | 2411 [1326, 3766] | 3167 [1667, 4688] | 3510 [2212, 5220] |
|  | (S)ame | 2764 [1468, 4400] | 2935 [1640, 4433] | 2680 [1973, 3513] | 2680 [1993, 3478] |
| (F)ixed | (F)ixed | 480 [36, 1225] | 309 [-141, 1018] | 360 [218, 528] | 188 [46, 345] |
|  | (L)imited | 1685 [541, 3130] | 1857 [723, 3198] | 1738 [558, 3277] | 1909 [700, 3453] |
|  | (S)ame | 1098 [425, 1923] | 1098 [379, 2038] | 2220 [1487, 3150] | 2049 [1337, 2976] |

#### 6.5.2 Comparison of model families

A comparison of individual models did not provide a clear picture of what factor, or combination of factors is most important to describe human data best. To more directly address which factor contributes most to a model’s success, we define model families, where each family is a subset of all models that share a particular level of a particular factor, regardless of their levels of other factors (van den Berg et al., 2014; Shen & Ma, 2019). For example, all seven observer models that use an optimal decision rule would be included in the (O)ptimal level of the decision rule factor, regardless of their individual encoding or inference schemes. Similar to Shen & Ma, 2019, we compute the approximate marginal likelihood for level *i* for factor *F*, *F_i_*. To calculate this value, we first marginalize over all of our tested models *M*:

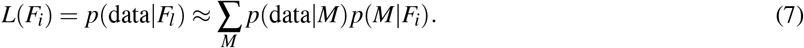

Next, we assume that all models containing level *i* of factor *F* are a priori equally probable, so that

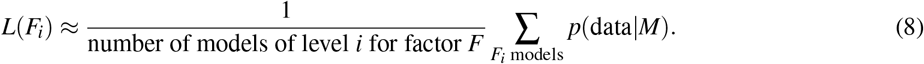

We approximate the log marginal likelihood of a given model with −0.5 * *AICc* (Burnham & Anderson, 2002):

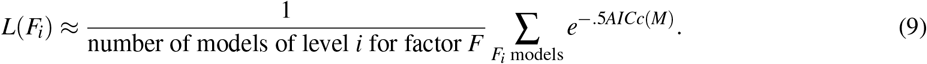

We define the *log level likelihood ratio* between level *i* and *j* as the ratio of their log marginal likelihoods:

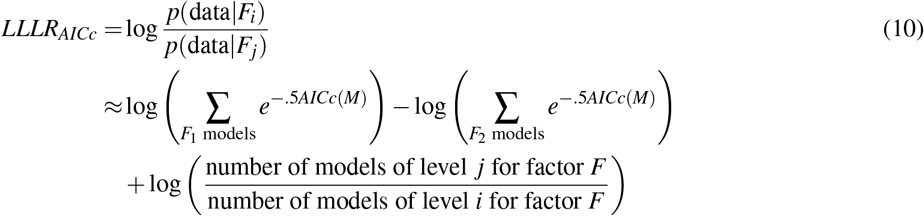

We also compute *LLLR*_BIC_, which approximates the log marginal likelihood of a given model with −0.5BIC, and more severely penalizes models with more parameters. To interpret the values for the LLLRs, we use Jeffrey’s scale, a common scale used when interpreting Bayes factors (Jeffreys, 1961).

##### Model factor 1: encoding scheme

The first model factor we explored is the observer’s encoding scheme. When using *LLLR*_AICc_, there is weak support that (V)ariable precision encoding outperforms (F)ixed precision encoding (summed [95% bootstrapped confidence interval (CI)] 84 [0, 181]). However, there is no evidence when using *LLLR*_BIC_ that either encoding scheme is favored (13 [−71, 116]). These results taken together imply weak and unreliable evidence in favor of a variable precision encoding.

##### Model factor 2: inference scheme

The second model factor is the observer’s inference scheme. As with the encoding scheme, (V)ariable precision fits better than (F)ixed when using AICc (105 [23, 200]), but not when using BIC (24 [−57, 126]). For both model comparison metrics, (L)imited and (S)ame inference schemes demonstrate a consistent lack of goodness of fit when compared to V (Limited – AICc: 500 [148, 928], BIC: 599 [253, 1047]; Same – AICc: 565 [204, 940], BIC: 565 [214, 974]) and F (Limited – AICc: 394 [68, 829], BIC: 575 [242, 996]) and Same: (AICc: 460 [124, 874], BIC 541, [193, 996]) inference schemes. L and S inference schemes perform similarly (AICc: 66 [26, 105], BIC: −34 [−85, 15]). These results provide strong evidence that participants use either a V or F inference scheme, but do not provide strong evidence to arbitrate between the two.

##### Model factor 3: decision rule

The third model factor is the observer’s decision rule. There is moderate evidence that the (O)ptimal decision rule fits better than the (M)ax decision rule when using BIC (69 [21, 132]) but not AICc (41 [−2, 108]). This result provides inconclusive evidence that participants are using the optimal decision rule.

##### Model factor 4: matching encoding and inference schemes

Finally, we define the fourth factor as whether encoding and inference schemes are matched (suggesting people have accurate representation of uncertainty) or mis-matched (suggesting people do not have accurate representation of uncertainty). While the previous three factors each investigate the effect of one model dimension on goodness-of-fit, this factor explores how the relationship between two model dimensions affect goodness-of-fit. This factor is arguably the most important aspect of the model in addressing our question of whether people maintain and use their uncertainty accurately over a working memory delay. The four models with matching encoding and inference schemes are VVO, VVM, FFO, and FFM, and the remaining 10 models are included in an “inference mismatched” level. There is very strong evidence that models where the inference and encoding schemes match fit data better than models that do not have encoding-matched inference schemes (AICc: 136 [51, 237], BIC: 200 [124, 291]). This result provides strong support that people accurately represent their uncertainty when completing the change detection task.

## 7 Discussion

In this paper, we investigated whether uncertainty is maintained and implicitly used in a working memory-based decision. First, we demonstrated through the Ellipse condition that people use uncertainty implicitly in a working memory task if that uncertainty information was available after the delay (i.e., if uncertainty did not need to be maintained). Second and more importantly, we showed through the Line condition that people not only use uncertainty, but maintain this information over the working memory delay. Finally, we factorially tested different model encoding schemes, inference schemes, and decision rules and found that people were indeed best described by models in which observers accurately maintain and use uncertainty in their decision.

First, we demonstrated through the Ellipse condition that people could use uncertainty implicitly in a working memory task if that uncertainty information was experimentally available. While the change detection task has been an experimental staple in the working memory literature (e.g., Luck & Vogel, 1997; Phillips, 1974; Pashler, 1988), the majority of these tasks feature large, categorical changes in the stimulus. In contrast, our task, which is a direct experimental replication of that of Keshvari and others (2012), featured changes that varied on a trial-to-trial basis. Trial-to-trial fluctuations in stimuli and withholding of feedback allow for a strongest test of probabilistic computation because observers would need to maintain a belief distribution over stimulus values to maximize performance in this task (Ma & Jazayeri, 2014). Through formal model comparison, we showed that all participants in the Ellipse condition are better fit by the Use Uncertainty model than the Ignore Uncertainty model. The Use Uncertainty model was identical to the model that was found to describe participant data best in the study by Keshvari et al. (2012). These results are also theoretically consistent with Devkar and others’ (2017) work, despite being slightly different tasks.

Second and more importantly, we showed through the Line condition that people not only use uncertainty, but maintain this information over the working memory delay. Like in the Ellipse condition, we found that all participants in the Line condition were better fit by the Use Uncertainty model than the Ignore Uncertainty model. However, the conclusion of this model comparison is critically different. In the Ellipse condition as well as in previous studies (Keshvari et al., 2012; Devkar et al., 2017), the ellipses were presented after the working memory delay, with the same reliability as before. With these experimental designs, reliability information could be used as a heuristic to inform uncertainty, thus not requiring this information to be maintained in memory. In other words, these previous studies cannot make any conclusions about the contents of working memory, only the decision-making process that follows it. Our result, in contrast, demonstrates that uncertainty was actually *maintained* in working memory, since the information was not available to the participants at the decision time through a heuristic such as ellipse reliability.

Finally, we conducted a factorial model comparison to investigate whether our conclusions were due to specific assumptions about model encoding schemes, inference schemes, and decision rules. When comparing individual models, models with different combinations of Variable or Fixed precision encoding scheme, Variable or Fixed precision inference scheme, and Optimal or Max decision rule were able to fit the data well. When comparing model families, we found that the only factor that clearly determined the goodness of fit of a model was whether the encoding and inference schemes were matched; only models with matching encoding and inference schemes captured human behavior qualitatively well, and these models were quantitatively superior to those without matching encoding and inference schemes. We thus conclude that the most important aspect of the model is that the observer accurately uses their uncertainty in the change detection decision, not the specifics of the encoding or inference process.

The results of this study corroborate those of previous studies, and extend them by providing evidence that people maintain uncertainty and use it *implicitly* and in a way that is *behaviorally-beneficial*. This is in contrast to studies that asked participants to make explicit reports such as confidence ratings (Rademaker et al., 2012; Vandenbroucke et al., 2014; Samaha & Postle, 2017), because use of uncertainty in these tasks are neither implicit nor behaviorally beneficial (i.e., your confidence rating doesnt affect your performance). Tasks such as the “choose best” (Fougnie et al., 2012; Suchow et al., 2017) and wager paradigms (Yoo et al., 2018; Honig et al., 2020) use uncertainty in a performance-relevant way, but it is arguable whether this use of uncertainty is implicit. These tasks can be considered implicit in the sense that there is a nontrivial mapping from uncertainty to performance-maximizing behavior in a post-perceptual decision, but explicit in the sense that this decision is related to a conscious feeling of trust in a memory. Conversely, a whole-report experiment by Adam and others (2017) analyzed by Schneegans and others (2020) clearly demonstrates an implicit use of uncertainty by showing participants reported remembered items in decreasing order of memory precision. However, unlike in our study, this use was not behaviorally beneficial; Adam and colleagues found a nonsignificant performance difference when allowing participants to freely report versus being probed on which items to report their memory of.

A typical, and reasonable, criticism of psychophysical experiments like the one presented in this paper is whether it can successfully distinguish whether people are representing uncertainty per se or some stimulus feature (i.e., ellipse reliability) as a *proxy* for it. Because observers with a Variable precision inference scheme represent uncertainty that fluctuates above and beyond stimulus variability, it seems unlikely that this observer would be representing uncertainty through stimulus reliability alone. This is, however, a valid criticism of the Fixed precision inference observer because the variability of their uncertainty representation fluctuates with stimulus reliability. While we did not directly test this alternative explanation (e.g., Barthelmé & Mamassian, 2010), we do not believe this criticism trivializes our results.

First, our results do not provide any evidence that people are simply maintaining ellipse reliability as a proxy for uncertainty. If stimulus reliability was used as a proxy for uncertainty, then models with a Fixed precision inference scheme would explain data best, independent of the encoding scheme. Instead, we found that the most important aspect of our models’ goodness of fit was that the encoding and inference schemes were *matched*, suggesting that accurate representation of uncertainty is the most important aspect to explaining human behavior.

Second, performing this task while maintaining a proxy to uncertainty is not as trivial as it may initially seem. Participants would still have to maintain this proxy to uncertainty over the working memory delay, then map this value to their decision rule in a way consistent with an optimal Bayesian observer. Since participants were not provided feedback on their performance, it is not obvious how they would have learned this mapping throughout the experiment. It is still possible that people do indeed map a stimulus feature to a decision in a way that is behaviorally and computationally indistinguishable from representing uncertainty itself. We do not believe this explanation would trivialize our results; either explanation still allows us to conclude that people maintain uncertainty (or a proxy of it) across a working memory delay, and use it implicitly in a task to improve performance.

Our results suggest that existing computational models of working memory that currently ignore uncertainty should be updated. For example, attractor network models currently maintain a point estimate of a single item feature through the mean of a stereotyped bump in a network of neurons (Ermentrout, 1998; Wang, 2001; Compte, 2006). Thus, there is typically no notion of uncertainty in this framework. Lim and Goldman (2014) demonstrated that altering the network connectivity and dynamics results in “negative-derivative feedback models,” in which networks can vary not only in mean but also in amplitude. Probabilistic population coding (PPC) and neural network models have implemented precision through input gain (Ma, Beck, Latham, & Pouget, 2006; Orhan & Ma, 2017). Additional research must investigate whether these negative-derivative feedback models can represent a memory’s precision through the amplitude of the network maintaining it, precision which could be read out from the observer as uncertainty.

Additionally, computational models could be used to decode uncertainty from neural activity in working memory tasks. Work in visual perception demonstrates that uncertainty information is represented in primary visual cortex (van Bergen, Ma, Pratte, & Jehee, 2015; van Bergen, 2019; Walker, Cotton, Ma, & Tolias, 2020; Hénaff, Boundy-Singer, Meding, Ziemba, & Goris, 2020). These studies built normative Bayesian models to infer stimulus value from fMRI BOLD signal. The likelihood of the stimulus, and thus uncertainty, could be read out from the models. Estimates of trial-specific uncertainty are positively correlated with error, suggesting that primary visual cortex held uncertainty information. Since working memories have been shown to be maintained in the same sensory areas with which they are perceived (e.g. Curtis & D’Esposito, 2003; Postle, 2006; D’Esposito & Postle, 2015; Harrison & Tong, 2009), perhaps visual working memory uncertainty is also stored in visual cortex. To more rigorously test the representation of uncertainty decoded from BOLD data, future studies can correlate decoded uncertainty with behavioral measures of uncertainty such as confidence ratings (Rademaker et al., 2012) or post-decision wagers (Yoo et al., 2018; Honig et al., 2020). Additionally, future studies can try to fit individual-trial data using these methods, which is more compelling evidence in favor of a model than a correlation.

Overall, this paper shows that people have uncertainty that reflects their memory noise at an item-specific level and they maintain this information over a working memory delay. This research demonstrates that there is other information, beyond a point estimate, maintained in working memory and used in later decisions.

## Acknowledgements

We thank Marissa Evans for the massive help collecting data for this study and Emin Orhan for collaborating on a previous iteration of this project. W.J.M is supported by award number R01EY020958. A.H.Y. was supported by training grant T32 EY7136-25. This work was supported in part through the NYU IT High Performance Computing resources, services, and staff expertise.

