## Supplementary materials for "Uncertainty is Maintained and Used in Working Memory"

### 8 Appendix

#### 8.1 Derivation of Decision Variable

In this section, we compute the decision variable from Eq. 1. Variables $\boldsymbol{\xi}$ and $\boldsymbol{\phi}$ denote the orientations of the stimuli on the first and second display, respectively, and each element in each is drawn from Uniform(0, $2\pi$). Variables $\boldsymbol{x}$ and $\boldsymbol{y}$ denote the memory for those orientations, respectively, and are drawn from von Mises distributions. $\Delta$ is the amount of orientation change and is drawn from Uniform(0, $2\pi$). $\boldsymbol{\Delta}$ is a vector of $\Delta$ on the $i^{\text{th}}$ location and 0s elsewhere. In order to compute the decision variable, we must marginalize over the unknown variables $\boldsymbol{\xi}, \boldsymbol{\phi}, \Delta$, and $\boldsymbol{\Delta}$.

$$
\begin{aligned}
p(\boldsymbol{x},\boldsymbol{y}|C)p(C) &= \iiiint p(\boldsymbol{x}|\boldsymbol{\xi})p(\boldsymbol{\xi})\, p(\boldsymbol{y}|\boldsymbol{\phi})p(\boldsymbol{\phi}|\boldsymbol{\Delta},\boldsymbol{\xi})p(\boldsymbol{\Delta}|C,\Delta)p(\Delta)p(C)d\boldsymbol{\xi}d\boldsymbol{\phi}d\boldsymbol{\Delta}d\Delta \\
&= p(C)\left(\frac{1}{2\pi}\right)^{N+1} \iiiint p(\boldsymbol{x}|\boldsymbol{\xi})p(\boldsymbol{y}|\boldsymbol{\phi})\delta(\boldsymbol{\phi}-(\boldsymbol{\xi}+\boldsymbol{\Delta})) \\
&\quad \left(\frac{1}{N}\sum_{i=1}^{N}\delta(\boldsymbol{\Delta}-C\Delta\mathbf{1}_i)\right) d\boldsymbol{\xi}d\boldsymbol{\phi}d\boldsymbol{\Delta}d\Delta \\
&= p(C)\left(\frac{1}{2\pi}\right)^{N+1} \frac{1}{N}\sum_{i=1}^{N}\iint p(\boldsymbol{x}|\boldsymbol{\xi})p(\boldsymbol{y}|\boldsymbol{\xi}+C\Delta\mathbf{1}_i)d\boldsymbol{\xi}d\Delta,
\end{aligned}
$$

where $\mathbf{1}_i$ is a vector of size $N$ with a 1 at the $i^{\text{th}}$ entry and 0s elsewhere. We then plug the above equation into the likelihood ratio:

$$
\begin{aligned}
\frac{p(\boldsymbol{x},\boldsymbol{y}|C=1)p(C=1)}{p(\boldsymbol{x},\boldsymbol{y}|C=0)p(C=0)} &= \frac{p(C=1)}{p(C=0)}\frac{\sum_{i=1}^{N}\iint p(\boldsymbol{x}|\boldsymbol{\xi})p(\boldsymbol{y}|\boldsymbol{\xi}+\Delta\mathbf{1}_i)d\boldsymbol{\xi}d\Delta}{\sum_{i=1}^{N}\iint p(\boldsymbol{x}|\boldsymbol{\xi})p(\boldsymbol{y}|\boldsymbol{\xi})d\boldsymbol{\xi}d\Delta} \\
&= \frac{p(C=1)}{p(C=0)}\frac{\sum_{i=1}^{N}\iint p(\boldsymbol{x}|\boldsymbol{\xi})p(\boldsymbol{y}|\boldsymbol{\xi}+\Delta\mathbf{1}_i)d\boldsymbol{\xi}d\Delta}{2\pi N\int p(\boldsymbol{x}|\boldsymbol{\xi})p(\boldsymbol{y}|\boldsymbol{\xi})d\boldsymbol{\xi}}
\end{aligned}
$$

Because the $N$ items are conditionally independent, we can break the expression up into a product of each item and further simplify.

$$
\begin{aligned}
d &= \frac{p(C=1)}{p(C=0)}\frac{\sum_{i=1}^{N}\int\prod_{j=1}^{N}\int p(x_j|\xi_j)p(y_j|\xi_j+\delta_{ij}\Delta)d\xi_j d\Delta}{2\pi N\prod_{j=1}^{N}\int p(x_j|\xi_j)p(y_j|\xi_j)d\xi_j} \\
&= \frac{p(C=1)}{p(C=0)}\sum_{i=1}^{N}\frac{\left(\prod_{j\neq i}\int p(x_j|\xi_j)p(y_j|\xi_j)d\xi_j\right)}{\left(\prod_{j\neq i}\int p(x_j|\xi_j)p(y_j|\xi_j)d\xi_j\right)}\frac{\int\left(\int p(x_i|\xi_i)p(y_i|\xi_i+\Delta)d\xi_i\right)d\Delta}{2\pi N\left(\int p(x_i|\xi_i)p(y_i|\xi_i)d\xi_i\right)} \\
&= \frac{p(C=1)}{p(C=0)}\sum_{i=1}^{N}\frac{\iint p(x_i|\xi_i)p(y_i|\xi_i+\Delta)d\xi_i d\Delta}{2\pi N\int p(x_i|\xi_i)p(y_i|\xi_i)d\xi_i} \\
&= \frac{p(C=1)}{p(C=0)}\sum_{i=1}^{N}\frac{\iint \mathrm{VM}(x_i;\xi_i,\kappa_{x,i})\mathrm{VM}(y_i;\xi_i+\Delta,\kappa_{y,i})d\xi_i d\Delta}{2\pi N\int \mathrm{VM}(x_i;\xi_i,\kappa_{x,i})\mathrm{VM}(y_i;\xi_i,\kappa_{y,i})d\xi_i} \\
&= \frac{p(C=1)}{p(C=0)}\sum_{i=1}^{N}\frac{\iint \mathrm{VM}(\xi_i;x_i,\kappa_{x,i})\mathrm{VM}(\Delta;y_i-\xi_i,\kappa_{y,i})d\xi_i d\Delta}{2\pi N\int \mathrm{VM}(x_i;\xi_i,\kappa_{x,i})\mathrm{VM}(y_i;\xi_i,\kappa_{y,i})d\xi_i} \\
&= \frac{p(C=1)}{p(C=0)}\sum_{i=1}^{N}\frac{1}{2\pi N\int \mathrm{VM}(x_i;\xi_i,\kappa_{x,i})\mathrm{VM}(y_i;\xi_i,\kappa_{y,i})d\xi_i} \\
&= \frac{p(C=1)}{p(C=0)}\sum_{i=1}^{N}\frac{1}{2\pi N\int \frac{I_0(\kappa)}{2\pi I_0(\kappa_{x,i})I_0(\kappa_{y,i})}\mathrm{VM}\left(\xi_i;\mu,\kappa\right)d\xi_i},
\end{aligned}
$$

where $\mu = x_i + \arctan(\sin(y_i - x_i), (\kappa_{x,i}/\kappa_{y,i}) + \cos(y_i - x_i))$ and $\kappa = \sqrt{\kappa_{x,i}^2 + \kappa_{y,i}^2 + 2\kappa_{x,i}\kappa_{y,i}\cos(x_i - y_i)}$.

$$= \frac{p(C=1)}{p(C=0)} \sum_{i=1}^{N} \frac{I_0(\kappa_{x,i}) I_0(\kappa_{y,i})}{N I_0(\kappa) \int \mathrm{VM}(\xi_i; \mu, \kappa)\, d\xi_i}$$
$$= \frac{p(C=1)}{p(C=0)} \frac{1}{N} \sum_{i=1}^{N} \frac{I_0(\kappa_{x,i}) I_0(\kappa_{y,i})}{I_0(\kappa)},$$

This produces the final expression of the decision variable. Eq. 2,

$$d = \frac{p(C=1)}{p(C=0)} \frac{1}{N} \sum_{i=1}^{N} d_i.$$

where

$$d_i = \frac{I_0(\kappa_{x,i}) I_0(\kappa_{y,i})}{I_0\left(\sqrt{\kappa_{x,i}^2 + \kappa_{y,i}^2 + 2\kappa_{x,i}\kappa_{y,i}\cos(x_i - y_i)}\right)}.$$

#### 8.2 Maximum-Likelihood Parameter Estimates

We report summary statistics of parameter estimates for only the Use Uncertainty (VVO) model in both conditions.

| | $\bar{J}_{\text{high}}$ | $\bar{J}_{\text{low}}$ | $\tau$ | $\sigma_{\text{d}}$ | $k$ | $\lambda$ |
|---|---|---|---|---|---|---|
| *M* | 33.45 | 12.39 | 30.16 | 0.66 | 4.87 | 0.09 |
| *SEM* | 4.32 | 3.32 | 8.53 | 0.30 | 1.73 | 0.02 |

Table 4: **Use Uncertainty (VVO) parameter estimates for the Ellipse condition**. Mean and SEM across participants for parameters in the model in which observers have variable precision (VP) encoding, assume VP encoding, and use an optimal decision rule.

| | $\bar{J}_{\text{high}}$ | $\bar{J}_{\text{low}}$ | $\bar{J}_{\text{line}}$ | $\tau$ | $\sigma_{\text{d}}$ | $k$ | $\lambda$ |
|---|---|---|---|---|---|---|---|
| *M* | 23.39 | 8.03 | 13.66 | 13.02 | 0.28 | 2.43 | 0.11 |
| *SEM* | 4.43 | 3.34 | 3.84 | 7.37 | 0.07 | 0.78 | 0.02 |

Table 5: **User Uncertainty (VVO) parameter estimates for the Line condition**. Mean and SEM across participants for parameters in the model in which observers have VP encoding, assume VP encoding, and use an optimal decision rule.

#### 8.3 Factorial Model Comparison: summed ΔAICc and ΔBIC

The sum of the ΔAICc and ΔBIC, along with their bootstrapped 95% CIs, for the Ellipse and Line conditions are displayed on 6 and Table 3, respectively. Summing the AICc and BIC explicitly assumes that participants are all fit by the same model.

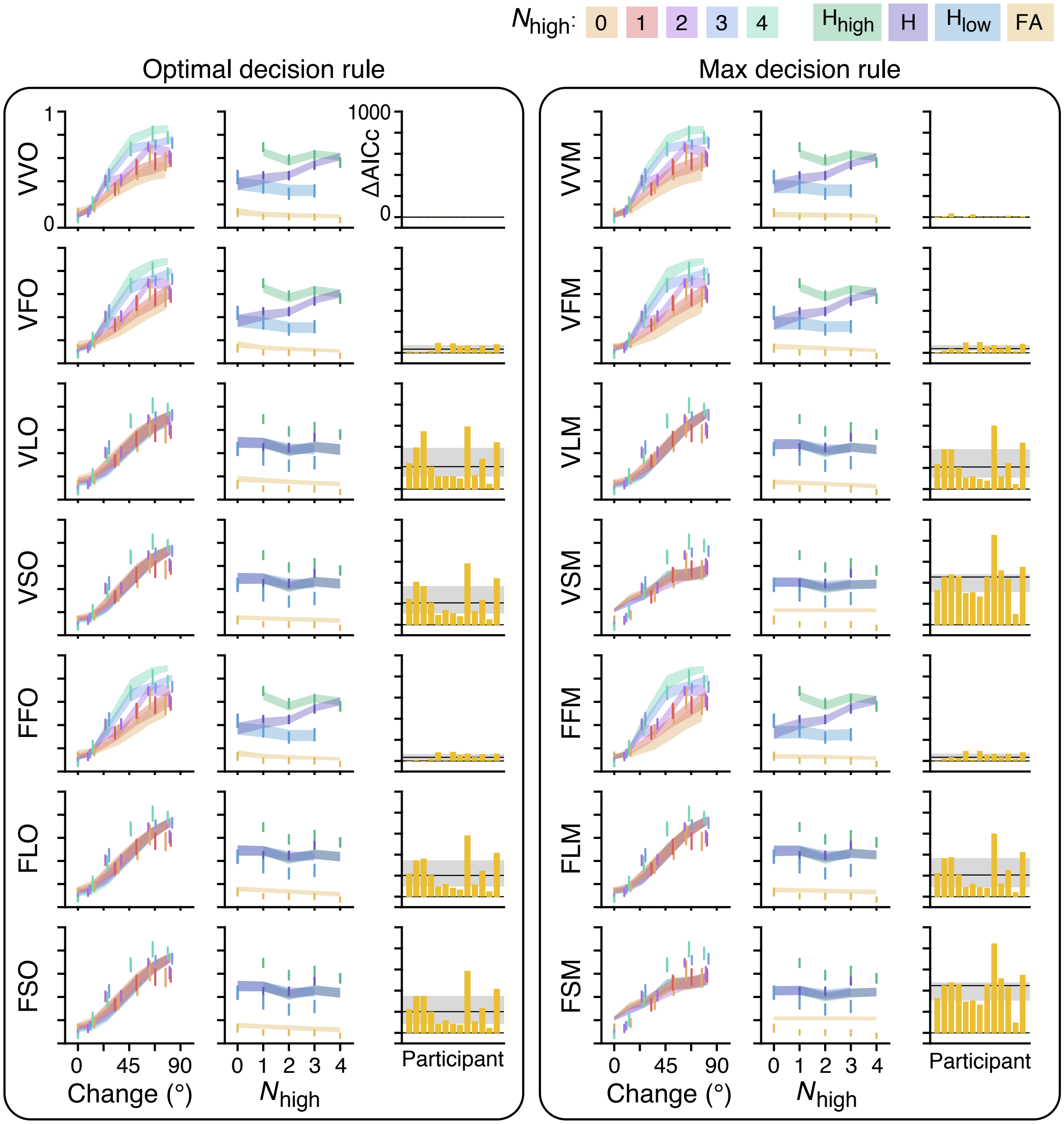


Figure 6: **Full model comparison: Ellipse condition.** Model predictions and performance of all possible combinations of different encoding, inference, and decision rules. $M \pm SEM$ data (error bars) and model fits (fills) for all models, organized into two columns by decision rule. For each model (each row within each column), the left graph illustrates the proportion report "change" as a function of amount of change. Color indicates the number of high reliability ellipses. The middle graph illustrates the proportion hits for high-reliability items (green), hits for low-reliability items (blue), hits averaged across the display (purple), and false alarms (yellow) as a function of number of high-reliability items. The right graph illustrates the individual participant ΔAICc, where positive numbers indicate VVO model is a better fit to the data. The grey horizontal line and grey shaded region illustrate the median and 95% bootstrapped confidence interval of the median across participants.

Both conditions suggest that the best model, when assuming all participants are from the same model, is VVO (Use Uncertainty) model, indicated by positive values. In the Ellipse condition, all sums and CIs are positive, suggesting the VVO model is superior. In the Line condition, the FFO model 95% CIs contains 0. Like in the main results, we cannot conclude from this metric that people's memory precision follows a Variable Precision memory model. However, there again is strong evidence that people accurately maintain their memory uncertainty.

| | | Decision Rule | | | |
|---|---|---|---|---|---|
| Encoding | Inference | (O)ptimal | | (M)ax | |
| | | ΔAICc | ΔBIC | ΔAICc | ΔBIC |
| (V)ariable | (V)ariable | 0 [0, 0] | 0 [0, 0] | 92 [10, 183] | 92 [9, 186] |
| | (F)ixed | 559 [330, 784] | 559 [336, 786] | 638 [408, 852] | 638 [421, 846] |
| | (L)imited | 3344 [2107, 4616] | 3516 [2303, 4693] | 3041 [1975, 4169] | 3213 [2128, 4355] |
| | (S)ame | 3091 [2015, 4321] | 3263 [2155, 4450] | 5532 [4318, 6796] | 5532 [4318, 6867] |
| (F)ixed | (F)ixed | 494 [255, 728] | 323 [101, 550] | 596 [378, 811] | 424 [199, 650] |
| | (L)imited | 2909 [1888, 4004] | 2909 [1894, 4025] | 2979 [1822, 4107] | 2979 [1969, 4212] |
| | (S)ame | 2895 [1871, 4164] | 2895 [1893, 3988] | 5504 [4322, 6765] | 5333 [4271, 6676] |

Table 6: **Summed ΔAICc and ΔBIC: Ellipse condition**. The sum and 95% bootstrapped confidence interval of the AICc and BIC differences between the optimal VVO (Use Uncertainty) model and others. A positive value indicates that the VVO model provides a better fit to the data. The cells corresponding to the Use (VVO) and Ignore (VSO) Uncertainty models are colored in blue.

#### 8.4 Group Bayesian Model Selection

We used group Bayesian Model Selection (BMS; Stephan, Penny, Daunizeau, Moran, & Friston, 2009) to examine whether it is reasonable to conclude that people use the same strategy in this task. BMS assumes that different participants may be best represented by different models, and the distribution of models is fixed but unknown across the population. It then infers the probability of each model from the log marginal likelihoods obtained for each model and participant. We used both $\frac{-\text{AICc}}{2}$ and $\frac{-\text{BIC}}{2}$ as approximations to the log marginal likelihoods. From this, we computed the protected exceedance probability, which is how likely a given model is to be more frequent than the other models in the comparison set, above and beyond chance. These values are presented in Table 7 and 8 for the Ellipse and Line condition, respectively.

In the Ellipse condition, the protected exceedance probabilities overwhelmingly suggest that participants correctly infer that they have variable precision encoding, and there is some evidence that most people use the optimal decision rule. For the Line condition, the two measures conflict about whether a VP or FP encoding and inference model best fits participants. In other words, it is unclear whether individual items are encoded (and correctly inferred) with noise above and beyond the noise generated from the different stimulus shapes. Nonetheless, both measures support the conclusion that people maintain an accurate representation of uncertainty over a working memory delay. The expected posterior probabilities are reported in Table 9 and 10 for Ellipse and Line condition, respectively. All models have some expected posterior frequency because of a Dirichlet uniform prior over models and a relatively low number of subjects to afford an informative random-effects analysis. Still, these results are consistent with our conclusion that people are aware of their memory variations, maintain it over a delay, and use it when making their decision.

| Encoding | Inference | Decision Rule | | | |
|---|---|---|---|---|---|
| | | (O)ptimal | | (M)ax | |
| | | $\frac{-\text{AICc}}{2}$ | $\frac{-\text{BIC}}{2}$ | $\frac{-\text{AICc}}{2}$ | $\frac{-\text{BIC}}{2}$ |
| (V)ariable | (V)ariable | 0.96 | 0.84 | 0.04 | 0.03 |
| | (F)ixed | 0.00 | 0.00 | 0.00 | 0.00 |
| | (L)imited | 0.00 | 0.00 | 0.00 | 0.00 |
| | (S)ame | 0.00 | 0.00 | 0.00 | 0.00 |
| (F)ixed | (F)ixed | 0.00 | 0.10 | 0.00 | 0.01 |
| | (L)imited | 0.00 | 0.00 | 0.00 | 0.00 |
| | (S)ame | 0.00 | 0.00 | 0.00 | 0.00 |

Table 7: **Protected exceedance probabilities: Ellipse condition**. Protected exceedance probabilities for each model, as estimated through group Bayesian model selection. The cells corresponding to the Use (VVO) and Ignore (VSO) Uncertainty models are colored in blue.

| Encoding | Inference | Decision Rule | | | |
|---|---|---|---|---|---|
| | | (O)ptimal | | (M)ax | |
| | | $\frac{-\text{AICc}}{2}$ | $\frac{-\text{BIC}}{2}$ | $\frac{-\text{AICc}}{2}$ | $\frac{-\text{BIC}}{2}$ |
| (V)ariable | (V)ariable | 0.78 | 0.01 | 0.12 | 0.04 |
| | (F)ixed | 0.01 | 0.00 | 0.00 | 0.00 |
| | (L)imited | 0.00 | 0.00 | 0.00 | 0.00 |
| | (S)ame | 0.00 | 0.00 | 0.00 | 0.00 |
| (F)ixed | (F)ixed | 0.03 | 0.93 | 0.00 | 0.00 |
| | (L)imited | 0.01 | 0.00 | 0.02 | 0.00 |
| | (S)ame | 0.01 | 0.00 | 0.00 | 0.00 |

Table 8: **Protected exceedance probabilities: Line condition**. Protected exceedance probabilities for each model, as estimated through group Bayesian model selection. The cells corresponding to the Use (VVO) and Ignore (VSO) Uncertainty models are colored in blue.

| Encoding | Inference | Decision Rule (O)ptimal | | (M)ax | |
|---|---|---|---|---|---|
| | | $\frac{-\text{AICc}}{2}$ | $\frac{-\text{BIC}}{2}$ | $\frac{-\text{AICc}}{2}$ | $\frac{-\text{BIC}}{2}$ |
| (V)ariable | (V)ariable | 0.39 | 0.30 | 0.15 | 0.10 |
| | (F)ixed | 0.04 | 0.04 | 0.04 | 0.04 |
| | (L)imited | 0.04 | 0.04 | 0.04 | 0.04 |
| | (S)ame | 0.04 | 0.04 | 0.04 | 0.04 |
| (F)ixed | (F)ixed | 0.05 | 0.15 | 0.04 | 0.07 |
| | (L)imited | 0.04 | 0.04 | 0.04 | 0.04 |
| | (S)ame | 0.04 | 0.04 | 0.04 | 0.04 |

Table 9: **Model posterior probabilities: Ellipse condition**. Expected posterior frequencies for each model, as estimated through group Bayesian model selection. The cells corresponding to the Use (VVO) and Ignore (VSO) Uncertainty models are colored in blue.

| Encoding | Inference | Decision Rule (O)ptimal | | (M)ax | |
|---|---|---|---|---|---|
| | | $\frac{-\text{AICc}}{2}$ | $\frac{-\text{BIC}}{2}$ | $\frac{-\text{AICc}}{2}$ | $\frac{-\text{BIC}}{2}$ |
| (V)ariable | (V)ariable | 0.28 | 0.09 | 0.15 | 0.13 |
| | (F)ixed | 0.04 | 0.04 | 0.04 | 0.04 |
| | (L)imited | 0.04 | 0.04 | 0.04 | 0.04 |
| | (S)ame | 0.04 | 0.04 | 0.04 | 0.04 |
| (F)ixed | (F)ixed | 0.09 | 0.34 | 0.04 | 0.04 |
| | (L)imited | 0.06 | 0.04 | 0.07 | 0.04 |
| | (S)ame | 0.05 | 0.06 | 0.04 | 0.04 |

Table 10: **Model posterior probabilities: Line condition**. Expected posterior frequencies for each model, as estimated through group Bayesian model selection. The cells corresponding to the Use (VVO) and Ignore (VSO) Uncertainty models are colored in blue.
